# Long-read transcriptome and other genomic resources for the angiosperm *Silene noctiflora*

**DOI:** 10.1101/2020.08.09.243378

**Authors:** Alissa M. Williams, Michael W. Itgen, Amanda K. Broz, Olivia G. Carter, Daniel B. Sloan

## Abstract

The angiosperm genus *Silene* is a model system for several traits of ecological and evolutionary significance in plants, including breeding system and sex chromosome evolution, host-pathogen interactions, invasive species biology, heavy metal tolerance, and cytonuclear interactions. Despite its importance, genomic resources for this large genus of approximately 850 species are scarce, with only one published whole-genome sequence (from the dioecious species *S. latifolia*). Here, we provide genomic and transcriptomic resources for a hermaphroditic representative of this genus (*S. noctiflora*), including a PacBio Iso-Seq transcriptome, which uses long-read, single-molecule sequencing technology to analyze full-length mRNA transcripts and identify paralogous genes and alternatively spliced genes. Using these data, we have assembled and annotated high-quality full-length cDNA sequences for approximately 17,000 *S. noctiflora* genes and 27,000 isoforms. We demonstrated the utility of these data to distinguish between recent and highly similar gene duplicates by identifying novel paralogous genes in an essential protease complex. Further, we provide a draft assembly for the approximately 2.7-Gb genome of this species, which is near the upper range of genome-size values reported for diploids in this genus and three-fold larger than the 0.9-Gb genome of *S. conica*, another species in the same subgenus. Karyotyping confirmed that *S. noctiflora* is a diploid, indicating that its large genome size is not due to polyploidization. These resources should facilitate further study and development of this genus as a model in plant ecology and evolution.

## Introduction

*Silene* is the largest genus in the angiosperm family Caryophyllaceae and serves as a model system in many fields of ecology and evolutionary biology (Bernasconi *et al.* 2009; Jafari *et al.* 2020). For instance, *Silene* is used to study breeding system evolution, as the genus includes hermaphroditic, gynodioecious, gynomonoecious, monoecious, and dioecious species (Desfeux *et al.* 1996; Charlesworth 2006). Gynodioecy (the coexistence of both hermaphroditic and male-sterile individuals) is thought to be the ancestral state of the genus (Desfeux *et al.* 1996) and is found in many extant *Silene* species as a result of cytoplasmic male sterility (CMS) factors (Taylor *et al.* 2001; Garraud *et al.* 2011). Dioecy, however, has evolved at least two times independently within *Silene*, including both ZW and XY sex determination systems (Mrackova *et al.* 2008; Slancarova *et al.* 2013; Balounova *et al.* 2019). Despite the diversity of *Silene* sexual systems, there is only one available whole genome sequence for the entire genus—from the dioecious species *S. latifolia*, which has heteromorphic XY sex chromosomes (Papadopulos *et al.* 2015; Krasovec *et al.* 2018). Whole genome resources are not available for any of the hermaphroditic species, which has limited comparative genomic studies into the evolution of dioecy within this genus. *Silene noctiflora* (**Figure 1**) is largely hermaphroditic but can produce a mixture of hermaphroditic and male-sterile flowers on the same plant (gynomonoecy) (Davis and Delph 2005). Also known as the night-flowering catchfly, this annual species is native to Eurasia and introduced throughout much of the world (McNeill 1980; Davis and Delph 2005).

**Figure 1:**
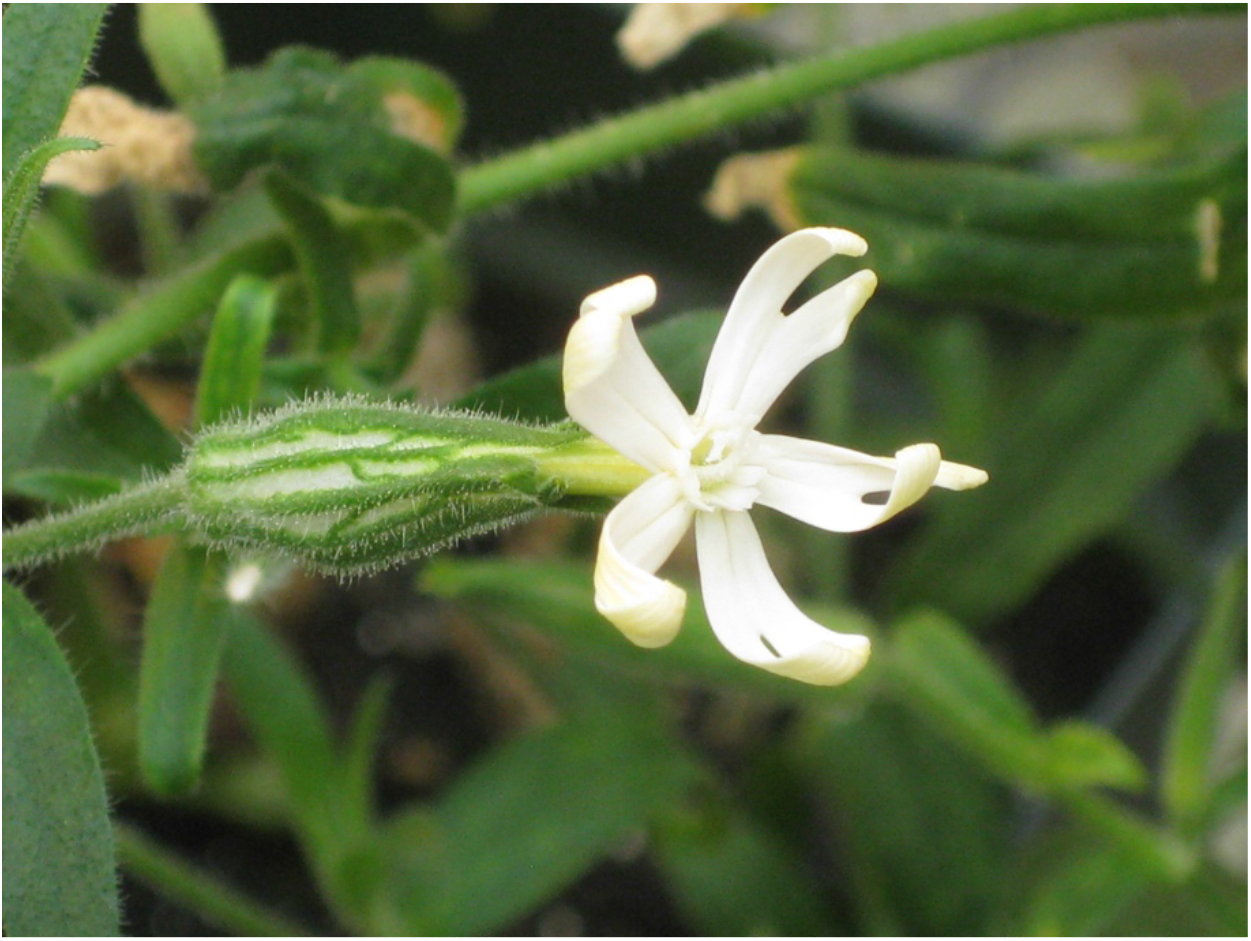
*Silene noctiflora*, also known as the night-flowering catchfly.

*Silene* is also used as a model system for investigating the coevolution between nuclear and cytoplasmic genomes (i.e., cytonuclear interactions), including in CMS systems (Olson and Mccauley 2002; Städler and Delph 2002; Klaas and Olson 2006; Garraud *et al.* 2011). In addition, there is considerable variation across the genus in organelle genome evolution. *Silene conica* and *S. noctiflora* have two of the largest known plant mitochondrial genomes at 11 Mb and 7 Mb, respectively (Sloan *et al.* 2012a). In contrast, the mitochondrial genome of *S. latifolia* is only 0.25 Mb, about 45 times smaller than that of *S. conica* (Sloan *et al.* 2012a). Interestingly, the *Silene* species with expanded mitogenomes also display unusually high evolutionary rates and stark structural changes in the mitochondrial genome (Mower *et al.* 2007; Sloan *et al.* 2012a). *Silene noctiflora*, for example, has a mitochondrial genome made up of around 60 circular-mapping chromosomes, and these chromosomes are rapidly gained and lost in different lineages (Wu and Sloan 2019). The plastid genomes in *S. conica* and *S. noctiflora* exhibit a correlated pattern of increased evolutionary rate—however, this pattern is found only in a subset of genes, and changes in plastid genome size and structure are more limited (Sloan *et al.* 2014). The natural variation in organelle genome evolution found in this genus has been used to study how these differences affect cytonuclear interactions (Havird *et al.* 2015; Williams *et al.* 2019).

The ability to use *Silene* as a model for cytonuclear evolution is still limited by the lack of extensive nuclear genome resources. Previous work has characterized *Silene* nuclear genome size and chromosome number. Nuclear genome sizes in the genus vary considerably, although not as starkly as mitochondrial genome sizes, ranging roughly 4.5-fold among diploids (haploid sizes of 0.71 to 3.23 Gb) and 8-fold when the tetraploid *S. stellata* (5.77 Gb) is included (Kruckeberg 1960; Siroký *et al.* 2001; Bai *et al.* 2012; Dagher-Kharrat *et al.* 2013; Pellicer and Leitch 2020). Most diploids in the genus, including *S. noctiflora*, have a chromosome number of 2n=24, which is likely the ancestral number (Bari 1973; McNeill 1980; Yildiz *et al.* 2008; Kemal *et al.* 2009; Gholipour and Sheidai 2010; Ghasemi *et al.* 2015; Mirzadeh Vaghefi and Jalili 2019). There are also numerous polyploid *Silene* species, including tetraploid, hexaploid, and octaploid forms (Kruckeberg 1960; Popp and Oxelman 2001, 2007; Popp *et al.* 2005; Bai *et al.* 2012). Most of the available nuclear sequence data comes from short-read RNA sequencing, which has been conducted on multiple *Silene* species (Blavet *et al.* 2011; Sloan *et al.* 2012b; Muyle *et al.* 2012; Casimiro-Soriguer *et al.* 2016; Havird *et al.* 2017; Bertrand *et al.* 2018; Balounova *et al.* 2019). These datasets have provided an important resource for molecular studies of *Silene*, but are limited because of the challenges associated with assembling short-read sequences, especially in distinguishing similar sequences arising from gene duplication, heterozygosity, and/or alternative splicing (Alkan *et al.* 2011; Schatz *et al.* 2012; Hahn *et al.* 2014; Lan *et al.* 2017).

Pacific Bioscience (PacBio) offers a long-read technology that involves sequencing single molecules, often leading to high error rates (Au *et al.* 2012; Rhoads and Au 2015; Hestand *et al.* 2016). However, these high error rates can be drastically reduced using circular consensus sequencing (CCS). CCS reads are generated by using hairpin adapters on each end of a double-stranded molecule, creating a circular, single-stranded topology (Wenger *et al.* 2019). This topology allows the polymerase to read the same full-length molecule multiple times over, generating an accurate consensus sequence (Ono *et al.* 2013; Wang *et al.* 2019). The application of PacBio CCS technology to reverse transcribed RNA (i.e., cDNA) samples is known as Iso-Seq and has been used to study the transcriptomes of many organisms, often in the context of identifying splice variants (Xu *et al.* 2015; Gordon *et al.* 2015; Rhoads and Au 2015; Guo *et al.* 2016; Abdel-Ghany *et al.* 2016; Wang *et al.* 2016; Weirather *et al.* 2017). Splice variants can be identified using CCS because this technology obtains consensus sequences for full-length single transcripts (Zhao *et al.* 2019). In the same way, CCS can also be used to distinguish paralogs or gene duplicates.

We have generated genomic resources critical for investigations into *S. noctiflora*, a species of interest due to its extremely unusual organelle evolution and resultant use as a model for cytonuclear interactions, as well as its status as a hermaphrodite in a genus representing many types of breeding system. We include a high-quality transcriptome using long-read PacBio Iso-Seq technology, genome size estimates, and a draft nuclear genome assembly. These resources will expand opportunities for molecular and ecological studies within the genus.

## Materials and Methods

### Plant growth conditions, tissue sampling, and nucleic acid extractions

Plants used for genome sequencing, Iso-Seq, and flow cytometry estimates of genome size were grown under standard greenhouse conditions with 16-hr light/8-hr dark at Colorado State University (**Table 1**). DNA for short-insert paired-end Illumina libraries was extracted from leaf tissue from a 7-week old *S. noctiflora* OPL individual using a Qiagen Plant DNeasy kit. Additional DNA was extracted from the same individual 6 weeks later using a modified CTAB protocol (Doyle and Doyle 1987) for construction of Illumina mate-pair libraries. For Iso-Seq library construction, RNA was extracted from a single 12-week old *S. noctiflora* OPL individual (grown from seed of the plant used for DNA extraction), using a Qiagen Plant RNeasy kit. RNA extractions were performed for four different tissue samples: 1) a large flower bud with calyx removed, 2) an entire smaller flower bud including calyx, 3) the most recent (top-most) pair of cauline leaves, and 4) one leaf from the second most recent pair of cauline leaves. The four RNA extractions were quantified with Qubit RNA BR kit (Thermo Fisher Scientific). Purity and integrity were assessed with a NanoDrop 2000 (Thermo Fisher Scientific) and TapeStation 2200 (Agilent Technologies).

**Table 1:** Genome sizes determined by flow cytometry.

| **Species** | **Population** | **Location** | **Samples (pg/2C)** | **Mean Genome Size** |  |
| --- | --- | --- | --- | --- | --- |
|  |  |  |  | **pg/2C** | **Gb/1C** |
| *Silene noctiflora* | OPL* | Opole, Poland | 5.65, 5.61, 5.46, 5.44 | 5.54 | 2.71 |
|  | OSR | Giles County, VA | 5.75, 5.61 | 5.68 | 2.78 |
|  | BRP | Nelson County, VA | 5.63, 5.57 | 5.60 | 2.74 |
| *Silene conica* | ABR | Abruzzo, Italy | 1.92, 1.92, 1.88 | 1.91 | 0.93 |
| *Silene vulgaris* | S9L | Giles County, VA | 2.19, 2.16 | 2.18 | 1.07 |
| *Silene latifolia* | UK2600 | Bedford County, VA | 5.46, 5.45 | 5.46 | 2.67 |

\*The *S. noctiflora* OPL population was used for Iso-Seq, genome assembly, and karyotyping

### PacBio Iso-Seq transcriptome sequencing and analysis

The four *S. noctiflora* RNA extractions (1.5 μg each) were pooled into a single sample and sent to the Arizona Genomics Institute for PacBio Iso-Seq library construction and sequencing. Library construction followed the standard PacBio Iso-Seq protocol (dated September 2018), and the library was sequenced with a PacBio Sequel (first generation) platform on two SMRT Cells.

Raw movie files of long-read, single-molecule sequences (one per SMRT cell) were processed using the PacBio Iso-Seq v3.1 pipeline (Anvar *et al.* 2018; Pacific Biosciences 2020). Circular consensus sequence calling was performed on each movie file separately using the command *ccs* with the recommended parameters *--noPolish* and *--minPasses 1.* Next, primer removal and demultiplexing was performed on each dataset by running the command *lima* with parameters *--isoseq* and *--no-pbi.* Poly(A) tails were trimmed and concatemers were removed using the *refine* command with the parameter *--require-polya.* Data from the two movies were merged at this point using the commands *dataset create --type TranscriptSet* and *dataset create --type SubreadSet.* Finally, the merged data were run through the *cluster* and *polish* commands.

Trinotate v3.2.0 (Bryant *et al.* 2017) was used to annotate the final polished sequences produced by the Iso-Seq pipeline. To complete this process, we used Transdecoder v5.5.0 (https://github.com/TransDecoder/TransDecoder/wiki), SQLite v3 (Kreibich 2010), NCBI BLAST + v2.2.29 (Camacho *et al.* 2009), HMMER v3.2.1 (including RNAMMER) (Lagesen *et al.* 2007; Potter *et al.* 2018), signalP v4 (Petersen *et al.* 2011), and tmhmm v2 (Krogh *et al.* 2001). The Pfam (Bateman *et al.* 2004) and UniProt (“UniProt” 2015) databases were included in the Trinotate installation. The transcripts and Transdecoder-predicted peptides were searched against the respective databases, following the standard Trinotate pipeline. All of these results were loaded into a Trinotate SQLite database.

Cogent v4.0.0 (https://github.com/Magdoll/Cogent/wiki), minimap2 v2.17 (Li 2018), and cDNA_Cupcake Py2 v8.7 (https://github.com/Magdoll/cDNA_Cupcake/wiki) were used to conduct family finding on the final sequences outputted by the Iso-Seq pipeline by partitioning sequences into gene families based on similarity. Next, coding genome reconstruction was performed on each gene family from the above step. Finally, a transcript-based genome was used to collapse redundant isoforms.

The Cogent family finding output was used with cDNA_Cupcake scripts (https://github.com/Magdoll/cDNA_Cupcake/wiki) to perform a rarefaction analysis (i.e. “collector’s curve”). First, a modified form of the script *make_file_for_sampling_from_collapsed.py* was run with the parameter *--include_single_exons* in order to include all transcripts in the analysis. Using the resultant file, *subsample.py* was run twice: once at the gene level (using the *pbgene* column) and once at the transcript level (using the *pbid* column). In both cases, parameters were kept as the default. Results of the rarefaction analysis were plotted in R using a modified version of the relevant cDNA_Cupcake script.

We used genes from the plastid caseinolytic protease (Clp) as a case study to assess the ability of Iso-Seq dataset to detect gene duplication events of various ages. To identify nuclear-encoded plastid Clp core genes in our dataset, we used blastn in conjunction with the Cogent family finding output. There are eight nuclear-encoded plastid Clp core genes in *Arabidopsis thaliana: CLPP3-6* and *CLPR1-4* (Nishimura and van Wijk 2015). Additionally, the genus *Silene* shares a duplication of *CLPP5*, denoted *CLPP5A* and *CLPP5B* (Rockenbach *et al.* 2016). We obtained the sequences of all nine of these genes from a previous study (Rockenbach *et al.* 2016) and used them as queries in blastn searches against the *S. noctiflora* Iso-Seq transcriptome. We then identified the BLAST hits in the Cogent output and based on those groups, we determined that eight of the nine nuclear-encoded Clp core subunits in *Silene* (including *CLPP5A* and *CLPP5B*) are single copy. However, in the case of *CLPR2*, two different Cogent families contained relevant transcripts, indicating a possible case of gene duplication. Sequence alignment of the transcripts within each Cogent family revealed that one family contained two unique sequences. These data, along with sequencing results from a separate cloning project, suggested that there are actually three distinct *CLPR2* sequences in *S. noctiflora*. We examined the other eight nuclear-encoded Clp gene Cogent families and found no evidence of additional duplications. In the subsequent phylogenetic analysis of *CLPR2*, we used the longest sequences from each of the three identified groups.

A phylogenetic tree was constructed using sequences from the three different *S. noctiflora CLPR2* genes. In addition to the three *S. noctiflora* sequences, we also included *Agrostemma githago, S. conica, S. latifolia, S. paradoxa*, and *S. vulgaris CLPR2* sequences from a previous study (Rockenbach *et al.* 2016), as well as three *S. undulata CLPR2* sequences identified using blastn against the *S. undulata* TSA database (accession GEYX00000000). All 11 sequences were aligned using the *einsi* option in MAFFT v7.222 (Katoh and Standley 2013), and trimmed at the 5’ end based on the trimming conducted in Rockenbach *et al.* (2016). The resultant sequence file was run through jModelTest v2.1.10 (Darriba *et al.* 2012) to choose a model of sequence evolution. We chose the top model based on the Bayesian Information Criterion (K80+I) and ran PhyML v3.3 (Guindon *et al.* 2010) with 1000 bootstrap replicates and 100 random starts.

### Genome size estimates by flow cytometry

Leaf or seedling samples were collected from multiple individuals of varying age (between 2 and 14 weeks) for each of our target *Silene* species and shipped fresh to Plant Cytometry Services (Schijndel, Netherlands). Genome sizes were determined using the CyStain PI Absolute P reagent kit (05-5502). Samples were chopped with a razor blade in 500 μl of ice-cold Extraction Buffer in a plastic petri dish, along with *Pachysandra terminalis* tissue as an internal standard (3.5 pg/2C). After 30-60 sec of incubation, 2 ml of Staining Buffer was added. Each sample was then passed through a nylon filter of 50 μm mesh size, and then incubated for 30+ min at room temperature. The filtered solution was then sent through a CyFlow ML flow cytometer (Partec GmbH). The fluorescence of the stained nuclei, which passed through the focus of a light beam with a 50 mW, 532 nm green laser, was measured by a photomultiplier and converted into voltage pulses. The voltage pulses were processed using Flomax version 2.4d (Partec) to yield integral and peak signals. Genome sizes were reported in units of pg/2C. The conversion used to report each size (x) in units of Gb was (x/2)*0.978 (Gregory *et al.* 2007).

### Karyotyping

*Silene noctiflora* OPL seeds were germinated on wet filter paper and grown for 5 days. Radicles were trimmed off and transferred to ice water for 24 hrs. The radicles were then fixed in a 3:1 solution of absolute ethanol and glacial acetic acid and stored at −20°C. Chromosomes were visualized using a squash preparation with Feulgen staining. Fixed radicles were rinsed in distilled water for 5 min at 20°C. Radicles were then hydrolyzed in 5M HCl at 20°C for 60 min followed by three rinses in distilled water. The hydrolyzed radicles were transferred to Schiff’s reagent to stain the DNA for 120 min at 20°C and were then destained by rinsing in SO2 water at 20°C three times for 2 min, two times for 10 min, once for 20 min, and then transferred to distilled water. Squashes were prepared by placing a piece of tissue in 45% acetic acid for 10 minutes and then minced on glass. A coverslip was placed over the minced tissue and pressed with enough pressure to produce a monolayer of nuclei. Slides were placed on dry ice for 1 min, and the coverslip was removed. The slides were transferred to 96% ethanol for 2 min, air dried, and mounted with mounting medium. Chromosomes were observed using a compound light microscope at 100× magnification.

### Genome sequencing and assembly

Extracted *S. noctiflora* OPL DNA samples were used for Illumina library construction and sequencing. A paired-end library with a target insert size of 275-bp was constructed at the Yale Center for Genome Analysis and sequenced on a 2×150-bp HiSeq 2500 run (three lanes). Two mate-pair libraries (with target insert sizes of 3-5 kb and 8-11 kb) were generated at GeneWiz and sequenced on a 2×150-bp HiSeq 2500 run (one lane each). Approximately 480M, 250M, and 230M read pairs were generated for the 275-bp, 3-5 kb, and 8-11 kb libraries, respectively. These reads are available via the NCBI SRA (accessions SRR9591157-SRR9591159). Reads were trimmed for quality and to remove 3’ adapters, using cutadapt v1.3 (Martin 2011) under the following paramters*: -n 3 -O 6 -q 20 -m 30 -a AGATCGGAAGAGCACACGTCTGAACTCCAGTCAC --paired-output.* The trimmed reads were assembled with ALLPATHS-LG release 44837 (Gnerre *et al.* 2011). Estimates of mean insert size and standard deviation for each library were provided as input for the assembly by first mapping a sample of reads to the published *S. noctiflora* plastid genome (GenBank accession JF715056.1). These estimates were as follows: 274 bp (± 22 bp), 3752 bp (± 419 bp), and 9873 bp (± 1283 bp).

### Data availability

The original subread bam files and final transcript sequences longer than 199 bp from the PacBio Iso-Seq transcriptome are available at NCBI Sequence Read Archive (SRA accession SRR11784995) and NCBI Transcriptome Shotgun Assembly Sequence Database (TSA accession GIOF01000000), respectively. The genome assembly has been deposited in GenBank (accession VHZZ00000000.1). Additional data have been provided at GitHub (https://github.com/alissawilliams/Silene_noctiflora_IsoSeq): 1) the full transcriptome as outputted by the PacBio Iso-Seq pipeline, 2) the annotation report for the transcriptome, 3) a custom script used to create a gene_trans_map file for our data in order to use Trinotate on non-Trinity-derived data, 4) the Cogent family finding output, and 5) the set of trimmed, aligned sequences used in the *CLPR2* phylogenetic analysis.

## Results and Discussion

### *Silene noctiflora* Iso-Seq transcriptome: Gene content and duplication

Sequencing of the Iso-Seq library on two Sequel SMRT Cells produced 711,625 and 686,576 reads for the first and second cells, respectively, where each read was derived from a single molecule. The two SMRT Cells differed substantially in data yield, with totals of 12,765,109 and 21,844,543 subreads, corresponding to subread counts of 17.9 and 31.8 per read, respectively. These reads were merged into 65,642 distinct high-quality transcripts according to the thresholds of the Iso-Seq 3.1 *merge* and *polish* commands. Of these transcripts, only 14 were found to be non-plant sequences, all of which were derived from *Frankliniella occidentalis* (the western flower thrip), a common greenhouse pest that likely contaminated our tissue samples.

We used the Cogent (https://github.com/Magdoll/Cogent/wiki) family finding algorithm to further collapse the 65,642 transcripts into 11,677 “gene families,” and then used the Cogent data along with Cupcake (https://github.com/Magdoll/cDNA_Cupcake/wiki) to conduct a rarefaction analysis. The rarefaction analysis, or “collector’s curve”, uses random sampling of reads to determine whether the extent of sequencing was sufficient to detect most of the genes and isoforms in our RNA sample. Based on this analysis, the Iso-Seq transcriptome contains 16,230 *S. noctiflora* genes and 27,860 isoforms (**Figure 2**). In both cases, the rarefaction analysis converged on a single estimate at 560,000 reads out of 594,988, indicating that we sequenced enough reads to essentially saturate our detection ability (which is also evident in the fact that the curves plateaued).

**Figure 2:**
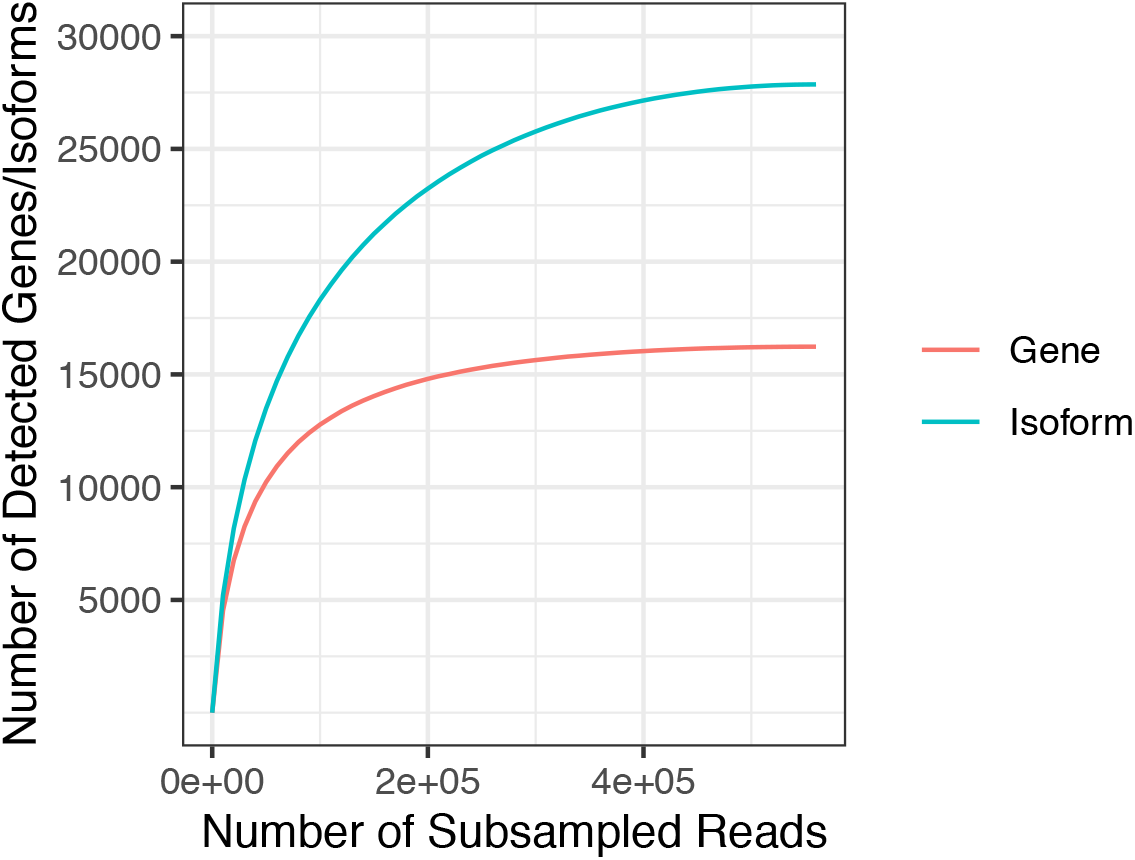
Rarefaction analysis of the *S. noctiflora* Iso-Seq transcriptome. Curves for both genes (red) and isoforms (blue) are depicted.

We wanted to test the ability of Iso-Seq to detect and distinguish paralogs of varying levels of divergence using the Cogent family finding output. To this end, we used a sample gene family— the core subunit genes of the plastid Clp complex, as they have a rich history of paralogy. In *E. coli* and most other bacteria, the core of the Clp complex, which is responsible for proteolysis, contains 14 identical subunits (Yu and Houry 2007). In cyanobacteria, gene duplication has led to four different core subunit-encoding genes (Stanne *et al.* 2007). Continued gene duplication in the land plant lineage has further reshaped this complex in plastids; the 14 core subunits are encoded by nine different genes in *A. thaliana*, eight of which are nuclear encoded (*CLPP3-6, CLPR1-4*), and one of which is plastid encoded (*clpP1*) (Nishimura and van Wijk 2015). Further, we had previously identified a more recent duplication of *CLPP5* in *Silene*, as well as duplications of the plastid-encoded *clpP1* in a small number of angiosperm species (Erixon and Oxelman 2008; Rockenbach *et al.* 2016; Williams *et al.* 2019).

We used the Cogent family finding output to examine the nine nuclear-encoded Clp core genes in *S. noctiflora.* The core genes *CLPP3, CLPP4, CLPP5A, CLPP5B, CLPP6, CLPR1, CLPR3*, and *CLPR4* were each represented by a single gene family in the Cogent output, whereas *clpR2* was represented by two gene families. Upon further examination, one of these families actually represented two different genes, yielding a total of three *CLPR2* genes in *S. noctiflora.* Thus, *CLPR2* was duplicated in this lineage, and then one paralog underwent a second gene duplication. Based on a phylogenetic analysis (**Figure 3**), these two duplications are shared with *S. undulata* but none of the other sampled *Silene* species. Thus, these duplications likely occurred after the *Silene* section Elisanthe (including *S. noctiflora*, *S. undulata*, and *S. turkestanica)* diverged from the other members of the genus (Jafari *et al.* 2020).

**Figure 3:**
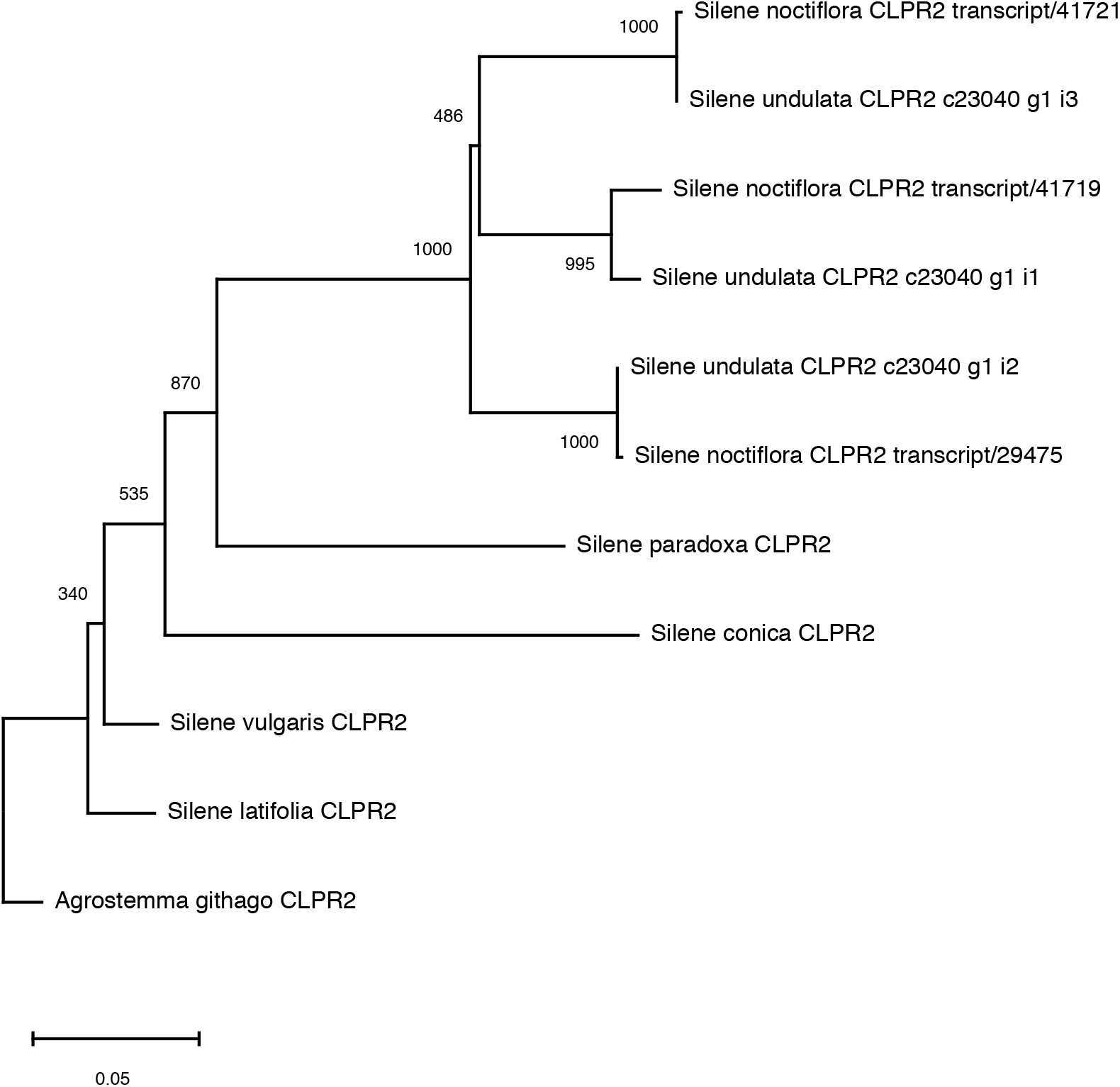
Phylogenetic analysis of *CLPR2* genes in *S. noctiflora* and related species. Branch lengths represent nucleotide sequence divergence. This tree was rooted on the *Agrostemma githago* sequence. The placement of *S. paradoxa* is in conflict with the species tree (Jafari *et al.* 2020), likely due to long branch attraction and the multiple independent evolutionary rate accelerations in this protein across *Silene* (Rockenbach *et al.* 2016).

The Iso-Seq data allowed us to identify transcripts from every known nuclear-encoded Clp core gene in *S. noctiflora*, including the closely related *CLPP5A* and *CLPP5B* subunits, as well as an additional, previously unreported triplication of *clpR2.* This result demonstrates that the Iso-Seq transcriptome provides highly accurate sequences, even for closely related paralogs that can be used in further study.

### *Silene* genome size estimates and chromosome number

Genome sizes of *S. noctiflora, S. conica, S. vulgaris*, and *S. latifolia* were determined using flow cytometry. Our estimates for *S. vulgaris* and *S. latifolia* (1.07 and 2.67 Gb, respectively; **Table 1**) were concordant with previously published estimates for these two species of 1.11 and 2.64 Gb (Costich *et al.* 1991; Siroký *et al.* 2001). Interestingly, despite their similar and extreme patterns of organelle evolution (Sloan *et al.* 2012a, 2014), including large mitochondrial genomes, *S. noctiflora* and *S. conica* have very different nuclear genome sizes. We found their respective genome sizes to be approximately 2.74 and 0.93 Gb, respectively (**Table 1**), which are on opposite ends of the spectrum for *Silene* diploids (Pellicer and Leitch 2020). The *S. noctiflora* nuclear genome is almost three-fold larger than that of *S. conica* suggesting that mitochondrial genome size is not necessarily correlated with nuclear genome size.

*S. noctiflora* has been previously reported as a diploid (2n=24) (McNeill 1980; Yildiz *et al.* 2008; Ghasemi *et al.* 2015). Given its relatively large genome size, we sought to confirm this result in our sampled population with a karyotype analysis (**Figure 4**), which indeed supported the conclusion that that *S. noctiflora* OPL is diploid.

**Figure 4:**
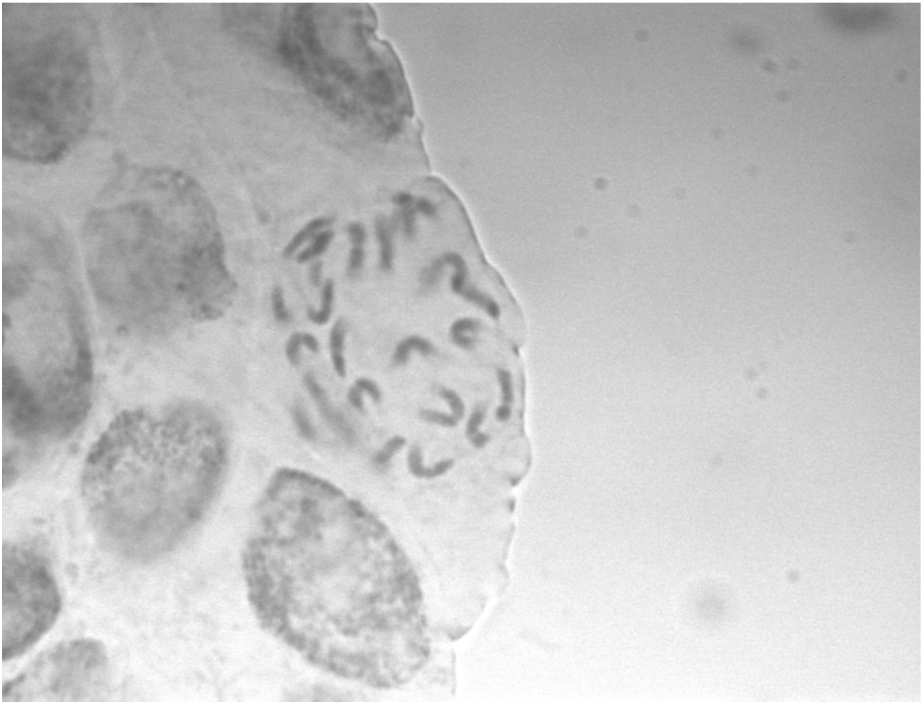
Micrograph verifying the diploidy of *Silene noctiflora* at 100× magnification.

### The *Silene noctiflora* nuclear genome

Illumina sequencing produced ~50× coverage of the *S. noctiflora* genome for a 275-bp paired-end library and ~15-20× for each of two mate-pair libraries. By performing a *de novo* assembly of these reads, we obtained a total assembly length (including estimated scaffold gaps) of 2.58 Gb, which is generally consistent with our estimate based on flow cytometry for *S. noctiflora* OPL (2.71 Gb). Given that we relied entirely on short-read sequencing technology, it was not surprising that the resulting assembly of this large genome was highly fragmented (79,768 scaffolds with a scaffold N50 of 59 kb). Moreover, assembly gaps made up 73% of the total scaffold length, presumably representing the highly repetitive content that is typical of plant nuclear genomes. As such, the assembled gap-free sequences amount to only about a quarter of the genome (702 Mb). This assembly should provide a useful resource to query for sequences of interest, especially in genic regions, and to compare against *S. latifolia* and other members of this genus. However, a more complete assembly that includes repetitive regions of the genome will require additional data from long-read technologies such as PacBio or nanopore sequencing.

## Acknowledgements

We thank Jocelyn Cuthbert and Zhiqiang Wu for assistance with plant growth and DNA extraction, Suzanne Royer for preliminary investigations into *Silene* karyotyping, and Joel Sharbrough for assistance with PacBio data analysis. This work was supported by a National Science Foundation (NSF) grant (MCB-1733227), start-up funds from Colorado State University, and graduate fellowships from NSF (DGE-1321845) and the National Institutes of Health (T32-GM132057).

